# Silylation of hydroxychloroquine enhances autophagy inhibition and antiproliferative activity in breast and pancreatic cancer cells

**DOI:** 10.1101/2025.11.26.690890

**Authors:** Michael J. Williams, Anuj K. Singh, Matthew D. Wittmer, Grant W. Anderson, Conor T. Ronayne, Venkatram R. Mereddy

## Abstract

**Introduction:** Autophagy is a key survival mechanism in cancer, and chloroquine (CQ) and hydroxychloroquine (HCQ) are widely used late-stage autophagy inhibitors. However, their low potency and the need for high doses limit antitumor efficacy and raise concerns about off-target toxicity. We therefore designed hydroxychloroquine silyl derivatives based on the hypothesis that increasing lipophilicity at the terminal 2-hydroxyethyl moiety enhances lysosomal accumulation and autophagy inhibition while preserving the 4-aminoquinoline pharmacophore.

**Methods:** Three HCQ silyl ethers-OTBS (**2**), OTIPS (**3**), and ODPS (**4**) have been synthesized by silylating the terminal hydroxyl of HCQ. Antiproliferative activity and autophagic flux along with apoptosis have been assessed for LC3B-II, p62/SQSTM1, and caspase-3-mediated PARP-1 cleavage.

**Results and Discussion:** The HCQ silyl derivatives retain the core pharmacophore and display improved or comparable antiproliferative activity relative to HCQ, achieving low-micromolar IC_50_ values. Candidate compounds **2** and **3** strongly induce LC3B-II and p62 accumulation than HCQ, consistent with enhanced blockade of autophagic flux. This autophagy disruption is accompanied by increased PARP-1 cleavage, particularly in aggressive 4T1 and MIAPaCa-2 cells, linking reinforced lysosomal/autophagy inhibition to apoptotic signaling.

**Conclusion:** These findings show that silylation of HCQ enhances autophagy inhibition and pro-apoptotic activity while maintaining pharmacology, identifying HCQ silyl derivatives as promising leads for autophagy-targeting antitumor agents.

## INTRODUCTION

Autophagy is a fundamental cellular process that maintains homeostasis by degrading and recycling damaged organelles and misfolded proteins **[1–3]**. This evolutionarily conserved catabolic mechanism provides eukaryotic cells with a sustainable source of biomolecules and energy, particularly under stressful conditions such as those encountered in the tumor microenvironment **[4–6]**. In cancer, however, autophagy exhibits a context-dependent and often paradoxical role. In early tumorigenesis, autophagy can act as a tumor suppressor by limiting oxidative stress, preserving genomic integrity, and preventing the accumulation of oncogenic damage. In contrast, in established tumors, autophagy frequently supports survival by promoting metabolic adaptation, therapy resistance, immune evasion, and metastasis **[1,2,4,7,8]**. This dual behaviour complicates therapeutic targeting but also highlights autophagy as an attractive, context-specific vulnerability in cancer **[5,6,9,10]**.

Autophagy is tightly regulated by a network of signalling pathways, including the mechanistic target of rapamycin kinase complex 1 (mTORC1), AMP-activated protein kinase (AMPK), unc-51-like autophagy activating kinase 1 (ULK1), and phosphatidylinositol 3-kinase (PI3K) complexes, which coordinate autophagosome initiation, nucleation, and maturation **[11–14]**. The microtubule-associated protein 1 light chain 3 (LC3) and sequestosome 1 (p62/SQSTM1) serve as key molecular markers and functional components of this pathway, regulating autophagosome formation, cargo selection, and subsequent delivery to lysosomes for degradation **[15,16]**. A detailed understanding of these regulatory mechanisms is essential for designing targeted strategies that either enhance or inhibit autophagy in a tumor- and context-specific manner **[1,2,5]**.

Pharmacological inhibition of autophagy disrupts this catabolic pathway, leading to the accumulation of dysfunctional organelles and misfolded proteins, increased cellular stress, and often reactive oxygen species (ROS) generation, ultimately driving cell death **[16–19]**. Blockade of autophagic flux, particularly at late stages, has emerged as a promising therapeutic approach to sensitize cancer cells to cell death, including apoptosis, and to enhance the efficacy of conventional and targeted therapies **[16,19–21]**. Antimalarial drugs chloroquine (CQ) and its derivative hydroxychloroquine (HCQ) are well-established lysosomotropic agents that primarily inhibit autophagy by increasing lysosomal pH and impairing autophagosome-lysosome fusion, thereby preventing degradation of sequestered cargo **[22–26]**. Similar late-stage autophagy inhibition is observed with other compounds such as Ginsenoside Ro and Liensinine, which also disrupt autophagosome-lysosome fusion and attenuate lysosomal cathepsin activity **[27,28]**. Through this mechanism, CQ and HCQ have been shown to resensitize cancer cells to diverse therapeutic agents across multiple malignancies, including lung, breast, and pancreatic cancers **[1,14,29,30]**. Despite their extensive clinical use, CQ and HCQ exhibit relatively low potency as autophagy inhibitors and require high doses to achieve sufficient antitumor effects, which can increase the risk of off-target toxicity **[29,31–33]**.

Several literature studies indicate that incorporation of silyl groups into biologically active molecules has excellent potential to improve pharmacology and pharmacokinetics along with reduction of side effects **[34–36]**. Incorporation of organosilicon motifs, including silyl ethers, generally increases molecular size and lipophilicity relative to the corresponding alcohols, which can enhance passive membrane permeability, tissue distribution, and in some cases oral bioavailability, while maintaining the core pharmacophore **[34,35]**. In addition, masking a polar hydroxyl as a silyl ether can reduce hydrogen bonding and phase-II conjugation, thereby improving metabolic stability and altering clearance profiles **[35,36]**. On this basis, we reasoned that converting the terminal hydroxyl of hydroxychloroquine into lipophilic silyl ethers could increase lipophilicity and membrane partitioning, thereby promoting lysosomal accumulation and potentially enhancing autophagy inhibition while preserving the 4-aminoquinoline pharmacophore. Accordingly, in this study we strategically incorporated and synthesized three hydroxychloroquine silyl ethers-hydroxychloroquine tert-butyldimethylsilyl, hydroxychloroquine triisopropylsilyl, and hydroxychloroquine tert-butyldiphenylsilyl, with the aim of improving lipophilicity and cellular uptake, thereby more effectively disrupt inhibit autophagy. Enhancing lysosomal delivery and autophagy blockade may translate into superior antiproliferative activity and improved chemosensitization in cancer cells.

## RESULTS AND DISCUSSION

### Design rationale and synthesis of hydroxychloroquine derivatives

CQ and HCQ, both historically used in the treatment of malaria showed potential to be used as autophagy inhibitors for cancer treatment **[32,33]**. The use of CQ or HCQ was associated with improvements in overall response rate and progression-free survival according to a meta-analysis of several clinical trials assessing their addition to standard chemo or radiation in various solid tumors **[37]**. Unfortunately, the improvement in overall survival from these trials has been limited due to CQ and HCQ’s low potency, insufficient suppression of autophagic flux at low concentrations, the requirement for high doses to achieve cancer cell kill, non-selective mechanisms, and intolerability at required high doses for anticancer effects **[38]**. These limitations highlight the need for development of new candidates with improved autophagy inhibition at low concentrations, high selectivity, and favorable druglike properties. We envisioned that incorporating silyl groups on to HCQ would improve its potency against cancer cells due to better lipophilic properties. We selected HCQ due to its ready availability, good clinical safety record as an antimalarial agent, and its autophagy inhibition properties in cancer cells. Moreover, the presence of a hydroxyl group in HCQ offers the introduction of silyl groups a straightforward process. We hypothesized that masking this hydroxyl in HCQ as a silyl ether would increase overall lipophilicity, enhance passive diffusion across cellular membranes, and promote lysosomotropic accumulation through improved membrane partitioning in combination with the basic side chain. Accordingly, HCQ **1** was transformed into three novel derivatives **2-4** via silylation of the terminal hydroxyl group **(Scheme 1)**. In a typical procedure, the hydroxyl group of HCQ acts as the nucleophile and reacts with the appropriate silyl chloride in the presence of base, affording hydroxychloroquine tert-butyldimethylsilyl ether (**2**), hydroxychloroquine triisopropylsilyl ether (**3**), and hydroxychloroquine tert-butyldiphenylsilyl ether (**4**). Introduction of these silyl groups markedly increases the steric bulk and hydrophobic surface area of the HCQ scaffold, changes expected to improve cellular uptake and membrane retention and to favour lysosomal accumulation, thereby amplifying the lysosomotropic behaviour that underlies autophagy inhibition by CQ/HCQ.

### Hydroxychloroquine derivatives (**2-4**) inhibit cancer cell proliferation in breast and pancreatic cancer cells

To evaluate the anticancer activity of HCQ **1** and its derivatives **2-4**, we assessed their effects on breast and pancreatic cancer cell viability in vitro. We used a panel of breast cancer cell lines comprising murine isogenic 67NR and 4T1 cells, along with the human triple-negative breast cancer line MDA-MB-231. In addition, we examined the human pancreatic ductal adenocarcinoma cell lines MIAPaCa-2 and BxPC-3. Using the 3-(4,5-dimethylthiazol-2-yl)-2,5-diphenyltetrazolium bromide (MTT) assay, we determined that HCQ **1** inhibited cancer cell proliferation with IC_50_ values ranging from 13-27µM in breast cancer cells and 22-44µM in the pancreatic cancer lines (**Table 1**). Interestingly, the silylated derivatives HCQ-OTBS (**2**), HCQ-OTIPS (**3**) and HCQ-ODPS (**4**) displayed low-micromolar activity, with IC_50_ values spanning approximately 0.67-7.65µM across the same panel (**Table 1**). These results illustrate that the antiproliferative activity of HCQ is largely conserved among derivatives **2-4** and, in several contexts, is enhanced, consistent with the predicted benefits of increased lipophilicity and better cellular permeabilization. Given that CQ/HCQ primarily act as late-stage autophagy inhibitors, the low-micromolar IC_50_ values observed here suggest that disruption of cytoprotective autophagy contributes substantially to the growth-inhibitory effects of these compounds, prompting further evaluation of their impact on autophagic flux in these models.

**Table 1.** IC_50_ values of synthesized compounds in 67NR, 4T1, MDA-MB-231, BxPC-3 and MIAPaCa-2 cell lines. Data represents the average ± SEM of three independent experiments (n=3).

| IC<sub>50</sub> (μM) values of synthesized compounds in different cancer cell lines | | | | | |
| --- | --- | --- | --- | --- | --- |
| Sl. No. | 67NR | 4T1 | MDA-MB-231 | BxPC-3 | MIA PaCa-2 |
| 1 | 13.28±1.07 | 27.0±4.05 | 25.05±1.09 | 44.23±2.6 | 22.2±3.1 |
| 2 | 0.679±0.05 | 3.59±3.13 | 2.67±1.01 | 2.54±1.08 | 3.06±1.53 |
| 3 | 1.88±1.24 | 5.47±2.54 | 2.88±0.81 | 2.75±0.79 | 3.29±1.37 |
| 4 | 4.11±1.70 | 7.75±3.21 | 30.41±10.2 | 22.43±2.9 | 17.73±4.9 |

Average ± SEM, minimum of three separate experimental results

### Mechanistic of Action for hydroxychloroquine derivatives using immunoblotting

To investigate the ability of the candidate compounds to inhibit autophagy, we examined the standard autophagy markers p62/SQSTM1 and LC3B-II in our panel of cancer cell lines. We further evaluated whether perturbation of autophagy correlated with induction of apoptosis by assessing caspase-3-mediated cleavage of PARP-1, a well-validated marker of apoptotic cell death. Immunoblotting was performed on 67NR, 4T1, and MIAPaCa-2 cells treated with DMSO, chloroquine (CQ), HCQ **1**, and the silylated HCQ derivatives HCQ-OTBS **2** and HCQ-OTIPS **3**. GAPDH was used as a loading control in all experiments (**Figures 1-3**). In non-metastatic 67NR cells (**Figure 1**), DMSO-treated controls showed low basal LC3B-II and modest p62 expression with minimal PARP-1 cleavage. CQ and HCQ **1** increased LC3B-II and caused detectable p62 accumulation, consistent with inhibition of autophagic flux, while **2** and **3** produced a further, albeit moderate, rise in p62, indicating more effective blockade of autophagosome clearance. PARP-1 cleavage remained weak in all treatments, suggesting that 67NR cells predominantly undergo cytostatic or non-apoptotic responses under these conditions. In the metastatic 4T1 counterpart (**Figure 2**), CQ and HCQ **1** induced robust LC3B-II accumulation and a clear increase in p62, and the silylated derivatives further intensified both signals, with **3** showing the strongest effect. PARP-1 cleavage was minimal in DMSO controls, modest with CQ and HCQ **1**, and most pronounced with **2** and **3**, linking enhanced autophagy blockade to increased apoptotic signaling. In MIAPaCa-2 pancreatic cells (**Figure 3**), CQ and HCQ **1** similarly induced strong LC3B-II and elevated p62, while HCQ derivatives produced comparable or greater LC3B-II, again most marked with **3**. All autophagy-inhibiting treatments triggered prominent PARP-1 cleavage, with CQ and **3** eliciting the highest apoptotic response.

**Figure 1.**
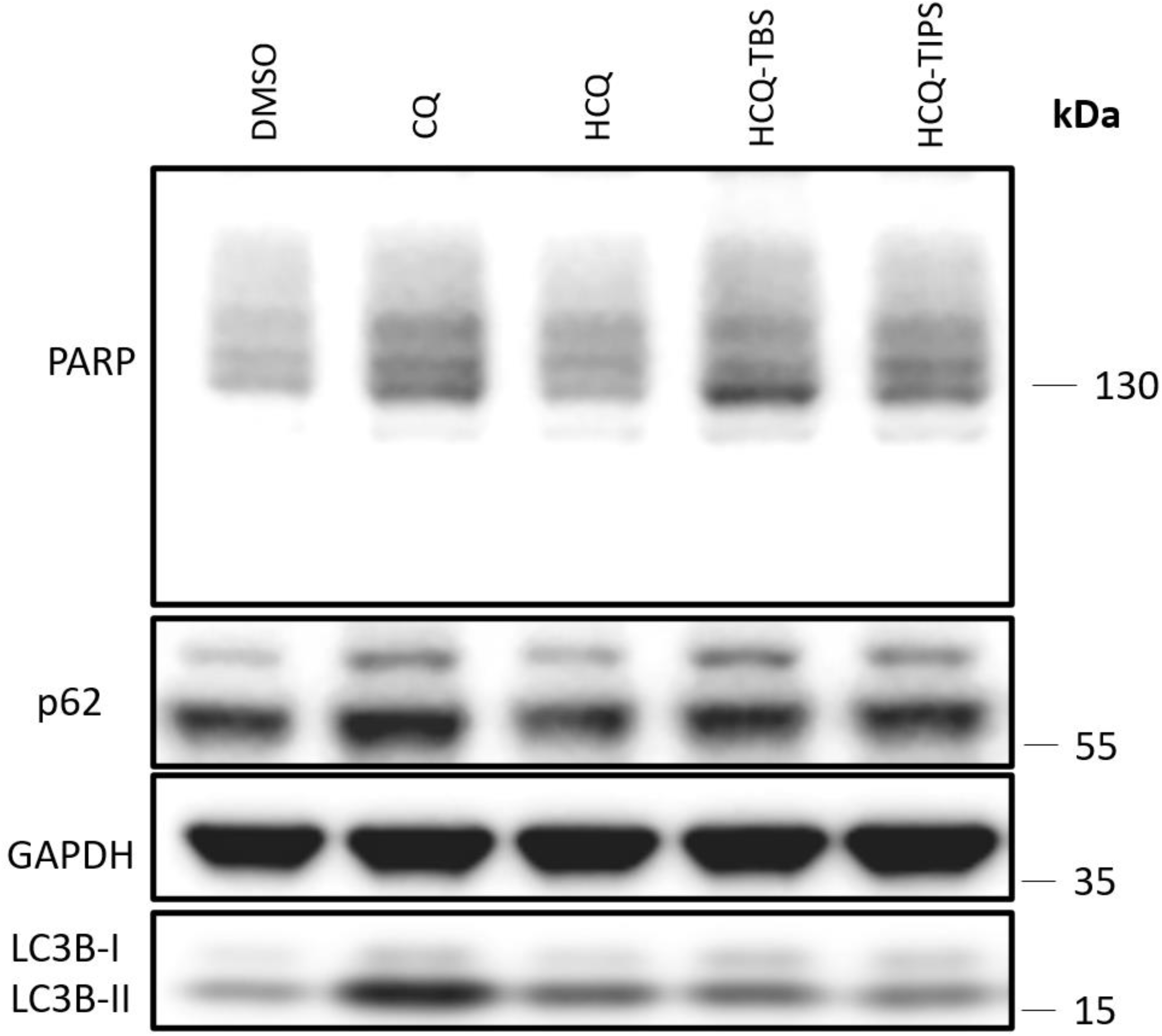
Western blot analysis of 67NR breast cancer cells treated for 24h with DMSO, CQ, HCQ, HCQ-TBS, or HCQ-TIPS. CQ and HCQ increase LC3B-II and p62 relative to vehicle, indicating inhibition of autophagic flux, while HCQ-TBS and HCQ-TIPS produce a modest further increase in p62 and LC3B-II. PARP cleavage remains minimal across treatments, suggesting predominantly non-apoptotic or cytostatic responses in 67NR cells at these doses. Blot shown is representative of at least three independent experiments.

**Figure 2.**
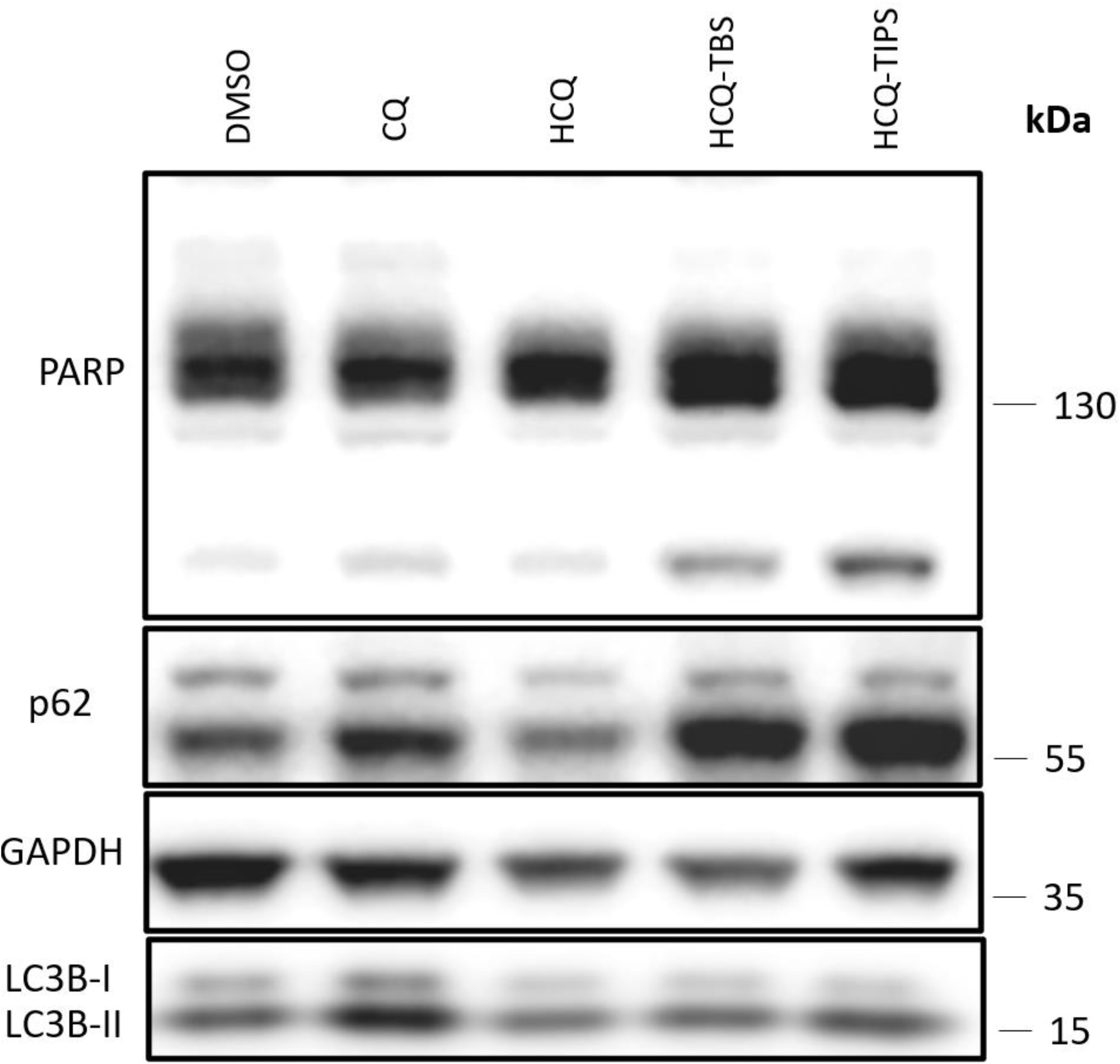
Western blot analysis of 4T1 breast cancer cells treated for 24h with DMSO, CQ, HCQ, HCQ-TBS, or HCQ-TIPS. HCQ and CQ increase LC3B-II and p62 relative to vehicle, indicating inhibition of autophagic flux, while HCQ-TBS and HCQ-TIPS further enhance LC3B-II and p62 accumulation. PARP cleavage is strongest with HCQ-TBS and HCQ-TIPS, linking the more robust autophagy blockade by the silyl derivatives to increased apoptotic signaling. Blot shown is representative of at least three independent experiments.

**Figure 3.**
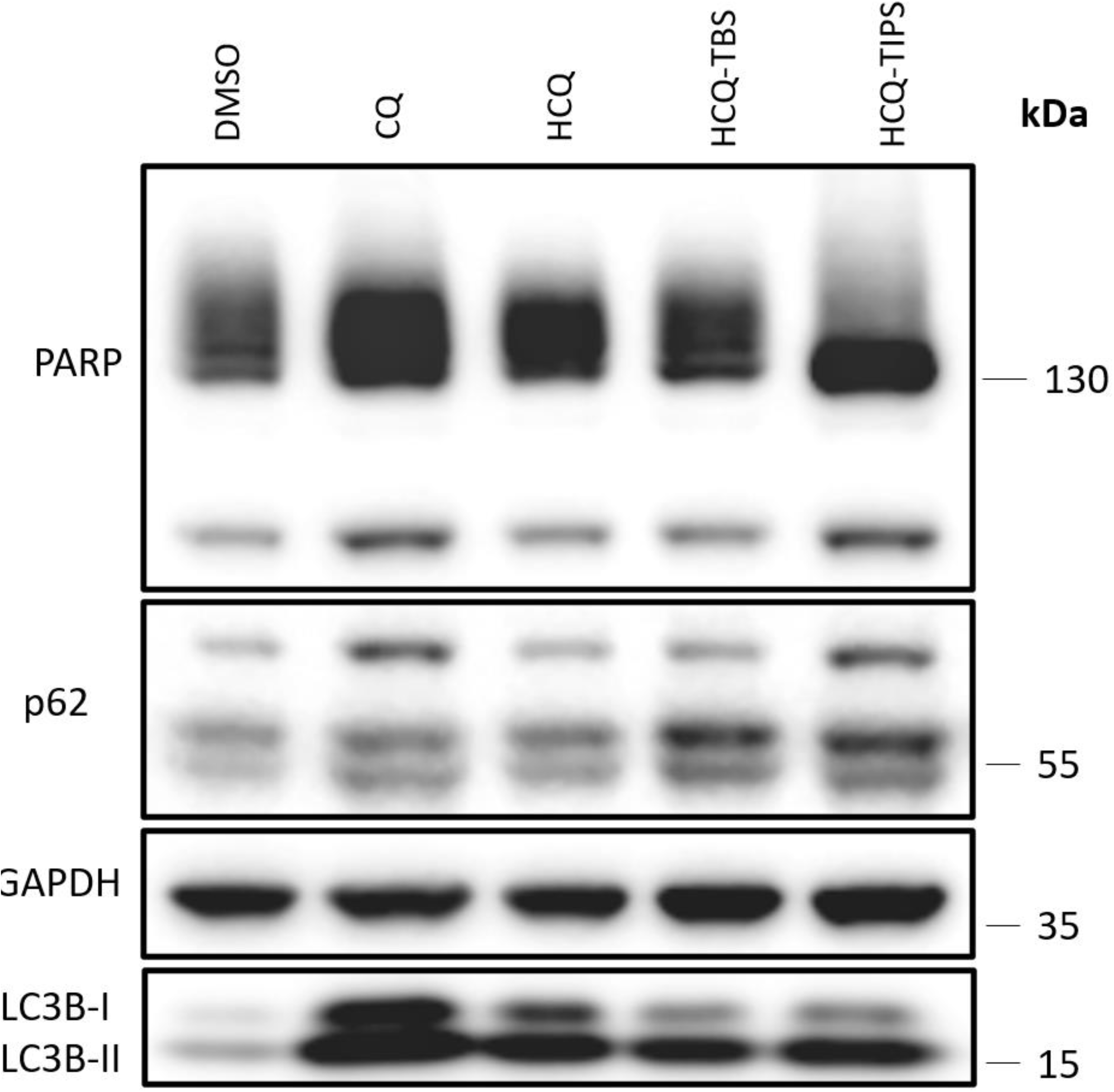
Western blot analysis of MIAPaCa-2 pancreatic cancer cells treated for 24h with DMSO, CQ, HCQ, HCQ-TBS, or HCQ-TIPS. All treatments increase LC3B-II and p62 relative to vehicle, indicating inhibition of autophagic flux, with HCQ-TIPS showing particularly strong LC3B-II and p62 accumulation. PARP cleavage is robust in response to CQ, HCQ, and especially HCQ-TIPS, linking enhanced autophagy blockade to pronounced apoptotic signaling in MIAPaCa-2 cells. Blot shown is representative of at least three independent experiments.

## CONCLUSION

In the current study, we designed and synthesized a series of hydroxychloroquine silyl derivatives based on the CQ/HCQ scaffold, guided by the hypothesis that increasing lipophilicity and steric bulk at the terminal 2-hydroxyethyl moiety would enhance lysosomal accumulation and autophagy inhibition. The resulting OTBS **2**, OTIPS **3** and ODPS **4** derivatives retained the core pharmacophore of HCQ and showed improved or comparable antiproliferative activity across a panel of breast (67NR, 4T1, MDA-MB-231) and pancreatic (BxPC-3, MIAPaCa-2) cancer cell lines, with low-micromolar IC_50_ values that in several cases exceeded those of the parent compound.

Mechanistic studies demonstrated that these derivatives effectively blocked autophagic flux, as evidenced by increased LC3B-II and p62 accumulation, and that this disruption was coupled to apoptotic signalling, reflected by caspase-3-mediated PARP-1 cleavage, particularly in aggressive 4T1 and MIAPaCa-2 cells. Together, our findings indicated that selective silylation of HCQ preserved autophagy-targeting activity while modestly reinforcing lysosomal blockade and pro-apoptotic effects. These HCQ derivatives therefore represent promising leads for the development of next-generation autophagy inhibitors and development as a potential antitumor agents for cancer therapy.

## METHODS

### Materials

Chloroquine diphosphate (CQ) and hydroxychloroquine sulfate (HCQ) were obtained from Ambeed (Arlington Heights, IL, USA). The silylation reagents tert-butyldimethylsilyl chloride (TBSCl), triisopropylsilyl chloride (TIPSCl), and tert-butyldiphenylsilyl chloride (TBDPSCl) were also purchased from Ambeed, and imidazole was obtained from Sigma-Aldrich (Merck, St. Louis, MO, USA). All other chemicals and solvents were of analytical grade and used as received unless otherwise specified.

Instrumentation:

– NMR: Varian Oxford-500; Bruker Ascend™ 400 (1H and 13C)
– HRMS (ESI): Bruker micrOTOF-Q III

Comprehensive characterization data for all new compounds are provided in the **Appendix**.

### Synthesis of N^1^-(2-((tert-butyldimethylsilyl)oxy)ethyl)-N^4^-(7-chloroquinolin-4-yl)-N^1^-ethylpentane-1,4-diamine (2)

2-((4-((7-chloroquinolin-4-yl)amino)pentyl)(ethyl)amino)ethan-1-ol (HCQ, 1; 1.2 mmol) was dissolved in dry dichloromethane (DCM, 15 mL) in a round-bottom flask, and imidazole (4.9 mmol) was added in one portion. The suspension was cooled to 0°C in an ice bath and stirred for 10 min. tert-Butyldimethylsilyl chloride (TBSCl, 1.5 mmol) was then added dropwise over 5 min, and the reaction mixture was allowed to warm to room temperature and stirred for 3-4 h. Progress was monitored by TLC (30% EtOAc/hexanes) until complete consumption of HCQ 1 was observed. The reaction was quenched with water and extracted with ethyl acetate (3X). Combined organic layers were washed thoroughly with water, dried over anhydrous MgSO_4_, filtered, and concentrated under reduced pressure. The residue was purified by column chromatography (10% EtOAc/hexanes, then DCM as needed) followed by recrystallization from hexanes to afford hydroxychloroquine tert-butyldimethylsilyl ether (2) as a solid in 80% yield.

### Synthesis of N^4^-(7-chloroquinolin-4-yl)-N^1^-ethyl-N^1^-(2-((triisopropylsilyl)oxy)ethyl)pentane-1,4-diamine (3)

Hydroxychloroquine triisopropylsilyl ether (3) was prepared using the same general procedure. HCQ 1 (1.2 mmol) was dissolved in dry DCM (15 mL), imidazole (4.9 mmol) was added, and the mixture was cooled to 0°C. Triisopropylsilyl chloride (TIPSCl, 1.5 mmol) was added dropwise over 5 min, and the reaction was stirred at room temperature for 5 h with TLC monitoring (25% EtOAc/hexanes). After completion, the mixture was quenched with water and extracted with ethyl acetate (3X). The combined organic extracts were washed with water, dried over anhydrous MgSO_4_, filtered, and concentrated. The crude material was purified by column chromatography (10% EtOAc/hexanes, then DCM) and recrystallized from hexanes to give compound 3 in 77% yield.

### Synthesis of N^1^-(2-((tert-butyldiphenylsilyl)oxy)ethyl)-N^4^-(7-chloroquinolin-4-yl)-N^1^-ethylpentane-1,4-diamine (4)

For compound 4, HCQ 1 (1.2 mmol) was dissolved in dry DCM (15 mL) and treated with imidazole (4.9 mmol) as above. The mixture was cooled to 0°C and tert-butyldiphenylsilyl chloride (TBDPSCl, 1.5 mmol) was added dropwise over 5 min. The reaction was allowed to warm to room temperature and stirred for 3-4 h, monitoring by TLC (30% EtOAc/hexanes) until starting material was consumed. The reaction was quenched with water and extracted with ethyl acetate (3X). Combined organic layers were washed vigorously with water to remove residual imidazole, dried over anhydrous MgSO_4_, filtered, and concentrated under reduced pressure. Purification by column chromatography (10% EtOAc/hexanes, then DCM) followed by recrystallization from hexanes afforded hydroxychloroquine tert-butyldiphenylsilyl ether (4) in 77% yield.

### Cell Culture

MDA-MB-231, 4T1, 67NR, MIAPaCa-2, and BxPC-3 cells (ATCC) were maintained at 37°C in a humidified incubator with 5% CO_2_. MDA-MB-231 cells were cultured in Dulbecco’s modified Eagle medium (DMEM) supplemented with 10% fetal bovine serum (FBS) and penicillin/streptomycin (50 U mL⁻¹ / 50 µg mL⁻¹). 4T1, 67NR, and BxPC-3 cells were maintained in RPMI-1640 medium containing 10% FBS and the same concentration of penicillin/streptomycin. MIAPaCa-2 cells were cultured in DMEM supplemented with 10% FBS, 2.5% horse serum, and penicillin/streptomycin (50 U mL⁻¹ / 50 µg mL⁻¹). Culture media were replaced 2-3 times per week, and cells were passaged at 70-80% confluence using standard trypsinization procedures. All cell lines were routinely monitored for morphology.

### Cell Proliferation Inhibition Assay (MTT)

Cell viability was assessed using the MTT colorimetric assay. Cells were seeded in 96-well plates at a density of 5x10³ cells per well in 100µL of complete growth medium and allowed to adhere overnight at 37°C in a humidified atmosphere of 5% CO₂. The following day, medium was replaced with fresh medium containing serial dilutions of each test compound; vehicle controls received an equivalent volume of DMSO. All conditions were set up in technical duplicate. Plates were incubated for 72h at 37°C, 5% CO₂. After treatment, MTT reagent was added to each well to a final concentration of 0.5mg/mL, and cells were incubated for an additional 4h to allow formation of formazan crystals. The medium was then removed, and the crystals were solubilized by adding 10% SDS in 0.01N HCl, followed by incubation for 4h at 37°C. Absorbance was measured at 570nm using a microplate reader. Background-corrected absorbance values were normalized to the mean of untreated control wells, which was set to 100% viability. Dose-response curves and 50% inhibitory concentrations (IC_50_) for each compound were generated using GraphPad Prism.

### Immunoblotting (Western Blot) Analysis of LC3B, p62, and PARP-1

Cancer cell lines 67NR, 4T1, and MIAPaCa-2 (ATCC) were seeded at 5.0x10⁵ cells per 60 mm dish and allowed to adhere for 24h at 37°C, 5% CO_2_. Cells were then treated with DMSO (vehicle), chloroquine, HCQ 1, or HCQ derivatives at 50% of their pre-determined IC_50_ values for 24h. Following treatment, media were aspirated, cells were washed twice with ice-cold PBS, and dishes were placed on ice and snap-frozen at -80°C until lysis. Cells were lysed in 1X RIPA buffer supplemented with protease and phosphatase inhibitors (50µL per plate), scraped, transferred to microcentrifuge tubes, briefly sonicated, and clarified by centrifugation at 4°C. Protein concentration was determined by BCA assay, and samples were mixed with 5X Laemmli sample buffer containing 10% β-mercaptoethanol. Equal amounts of protein (20µg) were separated on 4-12% SDS-PAGE gels in 1X MOPS running buffer and transferred to PVDF membranes.

Membranes were stained with Ponceau S to verify transfer, destained, and blocked in 5% non-fat dry milk in TBST for 1h at room temperature. Blots were incubated overnight at 4°C with primary antibodies against LC3B, p62/SQSTM1, PARP-1, and GAPDH (loading control) diluted in TBST containing 0.5% milk, washed, and then incubated with HRP-conjugated secondary antibodies for 1h at room temperature. After washing, signals were detected using enhanced chemiluminescence and imaged with a digital gel documentation system. Band intensities were quantified using ImageJ software.

## STATISTICAL ANALYSIS

All statistical analyses were performed using GraphPad Prism version 6.0 (GraphPad Software, San Diego, CA, USA). Data are presented as mean ± standard error of the mean (SEM) from at least three independent experiments, unless otherwise indicated in the figure legends. For comparisons of treatment groups to vehicle controls, one-way analysis of variance (ANOVA) followed by Dunnett’s multiple-comparisons test was used. Concentration–response data from viability assays were analysed by fitting nonlinear regression curves (log[inhibitor] vs. normalized response), from which 50% inhibitory concentrations (IC_50_) were derived. All graphing and curve fitting were performed in GraphPad Prism.

## STATEMENT FOR DATA SHARING AND DATA AVAILABILITY

The authors confirm that the data supporting the findings of this study are available within the article and its supplementary materials.

## AUTHORS’ CONTRIBUTIONS

XXXXXXXXXXXXXXXXXXXXXXXXX

## Scheme captions

**Scheme 1.**
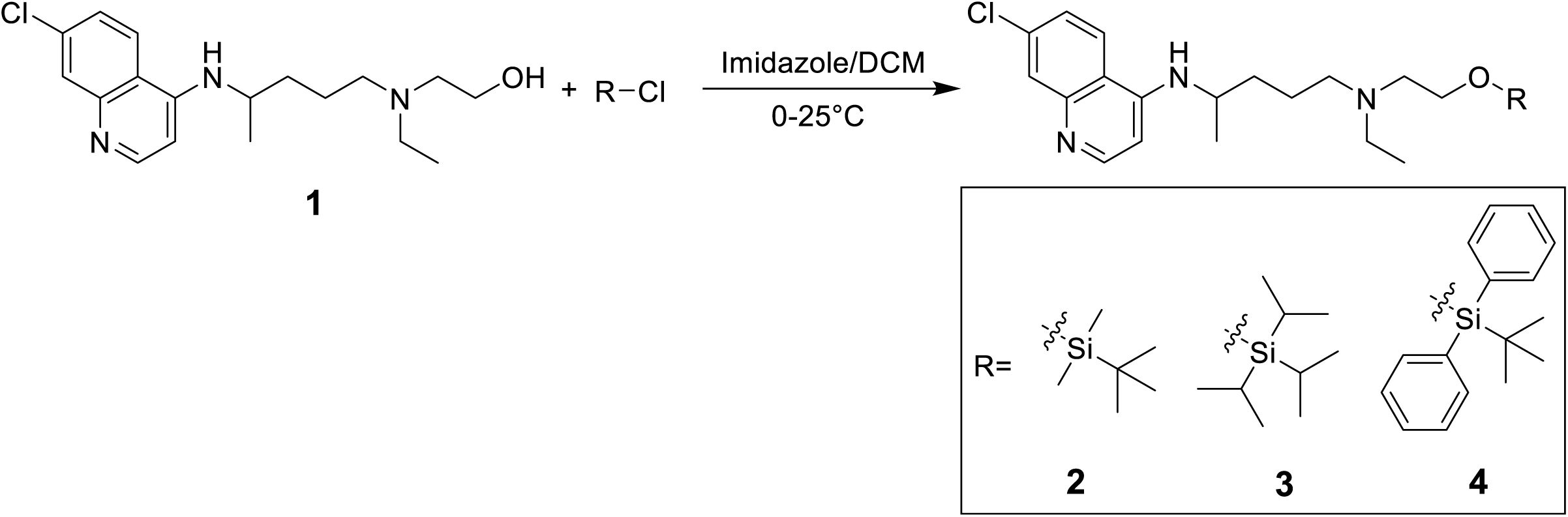
Synthesis of hydroxychloroquine silyl ethers **2-4** from parent hydroxychloroquine **1** which is treated with the appropriate chlorosilane (R-Cl) in the presence of imidazole in dry DCM at 0-25°C to afford the corresponding silyl ethers: tert-butyldimethylsilyl (OTBS, **2**), triisopropylsilyl (OTIPS, **3**), and tert-butyldiphenylsilyl (ODPS, **4**).

## APPENDIX

### Spectral Characterization

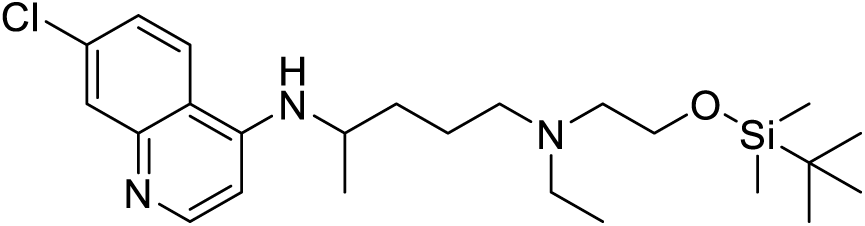
N^1^1-(2-((tert-butyldimethylsilyl)oxy)ethyl)-N4-(7-chloroquinolin-4-yl)-N1-ethylpentane-1,4-diamine

**^1^H NMR (400 MHz, Dimethyl Sulfoxide-d6):** δ 8.40 (d, J = 3.8 Hz, 1H), 8.38 (s, 1H), 7.8 (s, 1H), 7.44 (dd, 8.96 Hz, 1H), 6.94 (d, J = 7.96 Hz, 1H), 6.51 (d, J=5.44 Hz, 1H), 3.73 (q, J=12.76 Hz, 1H), 3.59 (t, J=12.76 Hz, 2H), 2.54-2.41 (m, 6H), 1.73-1.67 (m, 1H), 1.59-1.44 (m, 3H) 1.25 (d, J = 6.24 Hz, 3H), 0.95 (t, J=16.24 Hz, 3H), 0.87 (s, 9H), 0.00 (s, 6H)

**^13^C NMR (100 MHz, Dimethyl Sulfoxide-d6):** δ 152.3, 149.9, 149.8, 133.76, 127.89, 124.81, 124.18, 118.01, 99.2, 62.01, 55.65, 54.01, 48.1, 33.72, 26.20, 24.17, 20.31, 18.31, 12.43, -4.94

**HRMS (ESI) m/z:** Calculated for C_24_H_40_ClN_3_OSi+H^+^: 450.2602, found 450.2622

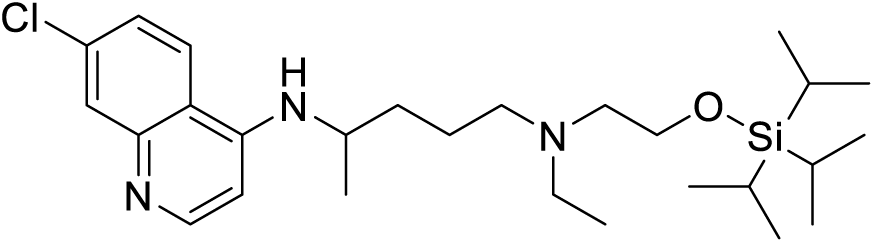
*N*^4^-(7-chloroquinolin-4-yl)-*N*^1^-ethyl-*N*^1^-(2-((triisopropylsilyl)oxy)ethyl)pentane-1,4-diamine

**^1^H NMR (400 MHz, Dimethyl Sulfoxide-d6):** δ 8.50 (d, J = 5.40 Hz, 1H), 7.95 (s, 1H), 7.66 (d, J = 2.08 Hz, 1H), 7.30 (m, 1H), 6.40 (d, J = 5.44 Hz, 1H), 5.10 (m, 1H), 3.75 (t, J = 6.68 Hz, 2H), 3.70 (m, 1H), 2.46-2.66 (m, 6H), 1.54-1.76 (m, 4H), 1.3 (d, J = 6.36 Hz, 3H), 0.95-1.11 (m, 25H)

**^13^C NMR (100 MHz, Dimethyl Sulfoxide-d6):** δ 152.0, 149.4, 148.9, 134.8, 128.9, 125.0, 121.0, 117.3, 99.3, 62.2, 55.7, 54.0, 48.5, 34.5, 24.2, 20.3, 18.0, 12.0, 11.7

**HRMS (ESI) m/z:** Calculated for C_21_H_31_F_3_N_2_OSi+H^+^: 492.2212, found 492.2213

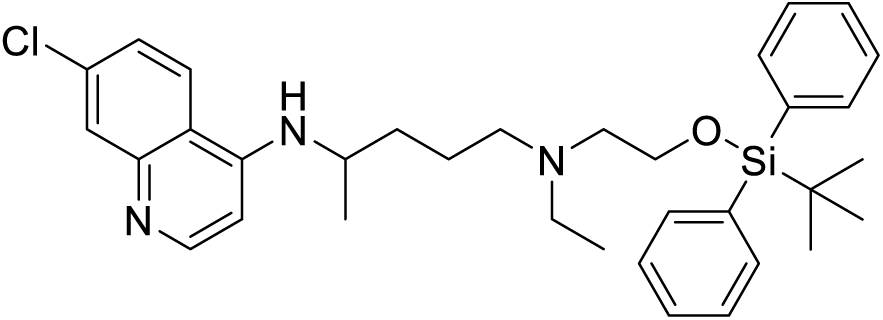
*N*^1^-(2-((*tert*-butyldiphenylsilyl)oxy)ethyl)-*N*^4^-(7-chloroquinolin-4-yl)-*N*^1^-ethylpentane-1,4-diamine

**^1^H NMR (400 MHz, Dimethyl Sulfoxide-d6):** δ Shift 8.50 (d, J = 5.40 Hz, 1H), 7.90 (s, 1H), 7.75-7.65 (m, 5H), 7.30-7.41 (m, 7H), 6.40 (d, J = 5.2 Hz, 1H), 4.90 (s, 1H), 3.66-3.75 (m, 1H), 3.60 (t, J = 5.32 Hz, 2H), 2.4-2.7 (m, 7H), 1.5-1.8 (m, 4H), 1.30 (d, J = 6.28 Hz, 3H), 1.10 (s, 9H), 1.01 (t, J = 7.00, 3H)

**^13^C NMR (100 MHz, Dimethyl Sulfoxide-d6):** δ Shift 152.0, 149.2, 149.0, 135.5, 134.9, 129.6, 128.8, 127.7, 125.2, 120.9, 117.2, 99.1, 58.3, 54.8, 53.0, 48.3, 47.5, 34.3, 26.6, 24.0, 20.4, 19.0, 11.6

**HRMS (ESI) m/z:** Calculated for C_21_H_31_F_3_N_2_OSi+H^+^: 574.2126, found 574.21647

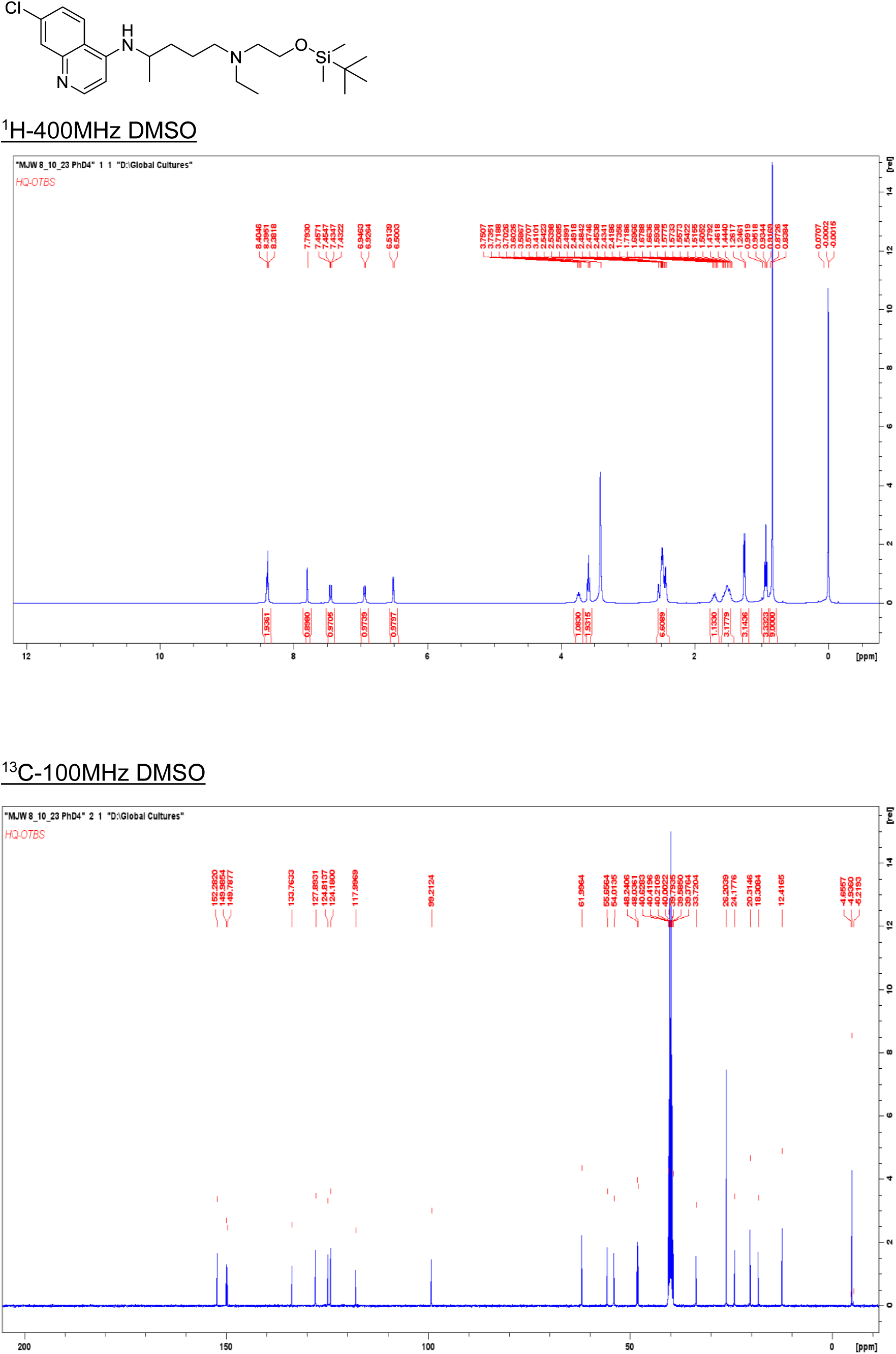

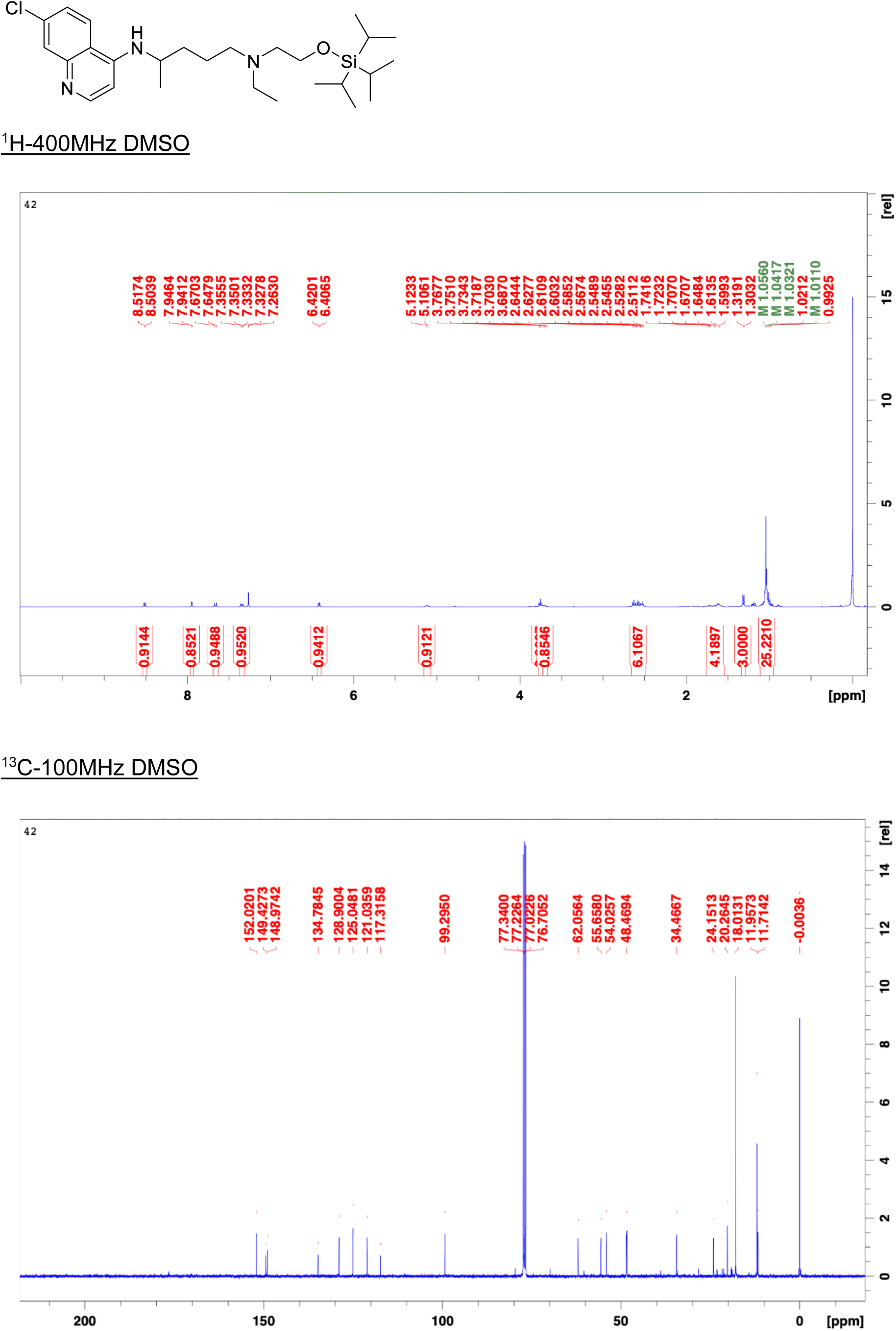

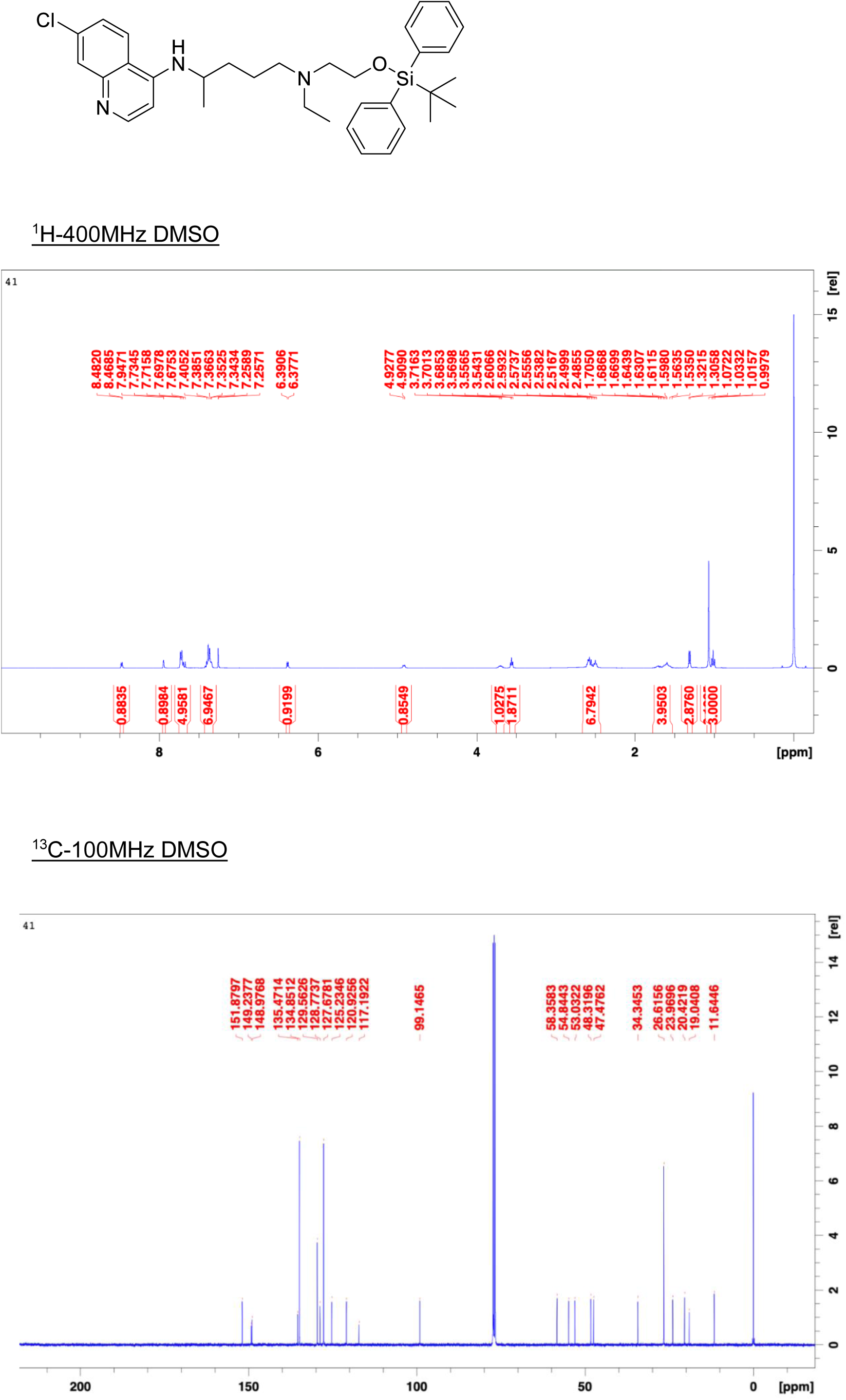

